# Why is the purse string not enough?

**DOI:** 10.64898/2026.08.05.743165

**Authors:** P. Vicente-Munuera, Jose J. Muñoz, Y. Mao

## Abstract

Wound repair is an important mechanism to preserve tissue integrity in organisms after injury. However, why different tissues exhibit different mechanisms to repair wounds is a long-standing question that remains unanswered. In this work, we theoretically explore the role of the purse string, an actomyosin contractile cable used by tissues to close small wounds. Does the tissue 3D geometry influence the efficiency of the purse string in driving wound closure? Using a 3D biophysical model, we study *in silico* tissues with the same cell volumes but different aspect ratios, ranging from squamous to thick and tall tissues. The model predicts that taller cells are easily deformed by the purse string. In contrast, very squamous cells require a very strong purse string that might demand additional cellular mechanisms to close the gap. These findings establish a theoretical framework to predict the optimal biophysical mechanisms of wound healing in different tissues.

**Graphical abstract:** Cells of different aspect ratios can be observed in a range of organisms with different function and mechanics. The wound healing efficiency of the purse string increases with the cell aspect ratio in our theoretical exploration.

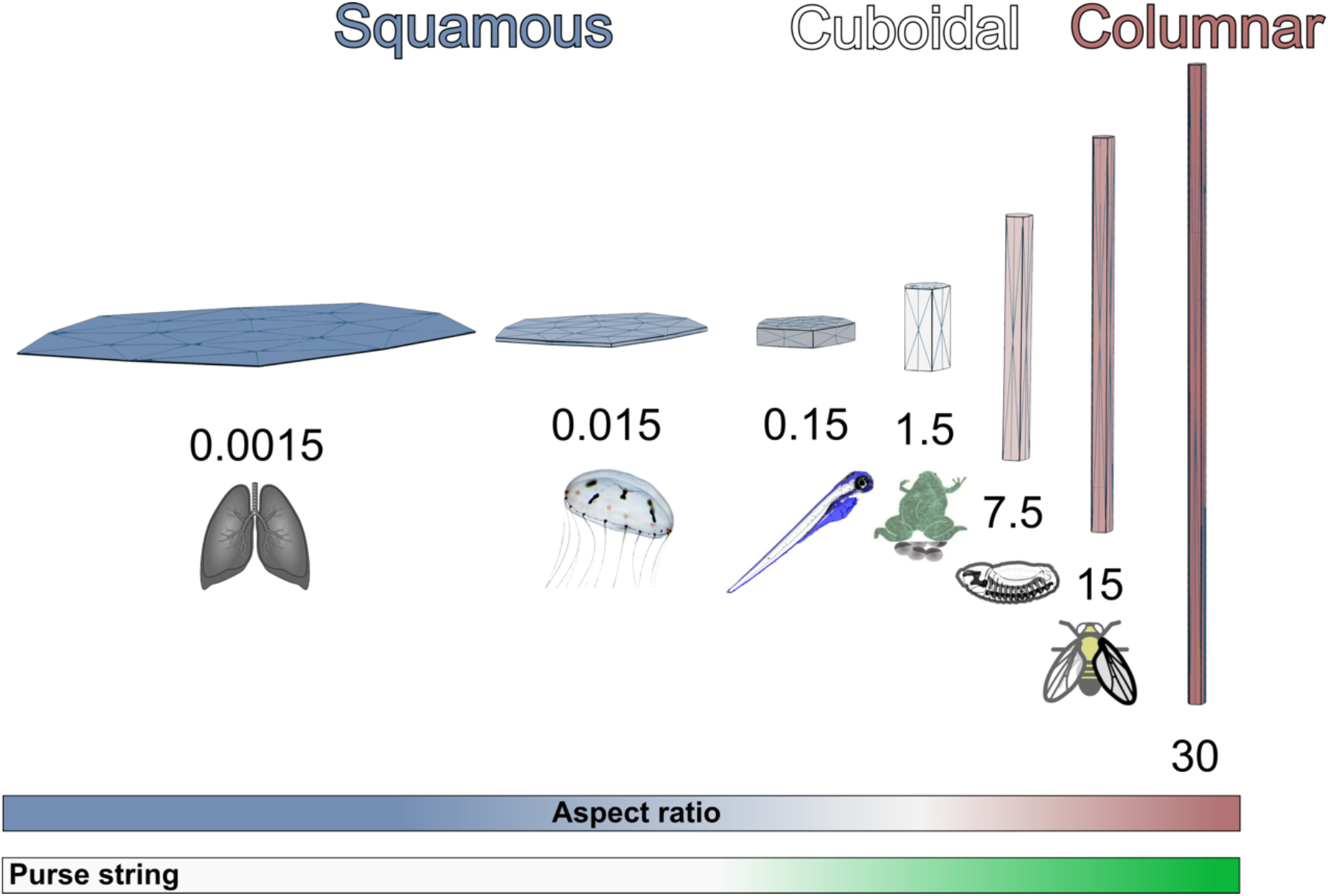

## Introduction

Epithelial tissues need to cope with mechanical insults that cause varying wound sizes. Large wounds typically require cell migration and proliferation to replace lost material^1–3^. In smaller wounds, however, tissues can recover without additional material or compensatory proliferation^4–6^. In these cases, the tissue mechanical environment becomes crucial, and cells have mastered different mechanisms depending on it.

Actomyosin-based mechanisms aid tissue healing by creating different contractile cables that line wound edge cells. A ring-like cable deforms the wound edge cells, reducing the wound size in the *XY* plane, like a so-called purse string^6–9^. Recent work has also shown straight lateral cables forming in the Z axis, shortening cells to aid wound closure^10^. How cells know where to recruit myosin II is still unknown. Still, the efficacy of the actomyosin purse string is very much dependent on the resistance of the cells to deformation forces and how much myosin II can be recruited to specific domains to generate such deformation forces. The resting surface tension determines the force the contractile cable must overcome to push forward and close the gap. If the resistance, surface or line tension is weak, a weak purse string would suffice to surpass the resting forces. This is thought to be related also to a fluid tissue^11,12^ where cells are softer and more deformable. In contrast, rigid and tense tissues would require stronger actomyosin mechanisms to overcome the resting tension. Given that the amount of contractile machinery that can be recruited is limited^13^, tissues might need to combine different mechanisms in a coordinated manner to ensure timely and efficient wound healing. Among others, tissues have shown a repertoire of additional mechanisms to close a gap: cell-cell intercalation, cell fusion, endoreplication, or protrusive lamellipodia^12^. Some studies have explored the efficiency of gap closure in squamous to cuboidal MDCK cells^14^. Still, it is yet to be understood how the efficacy of the contractile mechanism varies with cell and tissue shape.

The shape of any object is dictated by the forces exerted on it and its physical properties. In the context of the cell shape, the cell’s own material properties and the cell’s mechanical environment define its architecture and aspect ratio^15,16^. On one hand, the spatial organisation of myosin II, and the strength of cell-cell and cell-substrate adhesions can bias whether a cell would elongate, shrink or grow isotropically^17,18^. For instance, columnar cells were shown to have stronger cell-cell adhesion than squamous cells, meaning that the shared lateral face between two cells would be energetically linked to grow^19^. This suggests that there is a tight balance between apical and basal contractile forces, and cell-cell adhesion to maintain tissue geometry^20^. Similarly, for a cell to grow from squamous to cuboidal, assuming volume conservation, it can increase its apical and basal contractile forces to reduce its apical and basal area^18^. On the other hand, the tissue mechanical environment also impacts cell structure and function^21^. Previous works have shown how an elastic substrate can affect the shape and function of human mesenchymal stem cells^22^, and matrices’ viscoelasticity was found to define tissue spatial organisation^23^. Nonetheless, tissues have also adapted their wound healing mechanics depending on substrate stiffness^14,24,25^, suggesting a tight connection between cell shape, mechanics, and repair efficacy^12^. Still, changing the material properties of the cells while independently comparing different cell shapes *in vivo* is challenging, as biological systems are complex and interlinked.

Given the pleiotropy of performing such wound healing experiments in a biological setting, we use our recently developed computational 3D vertex model to analyse wounds in tissues with different cell aspect ratios. Others have utilised different 3D mechanical models to identify important aspects of wound healing^26,27^, while other studies have focused on understanding how tissues evolve and have different shapes^28–30^. Recently, we demonstrated that we could recapitulate *Drosophila* wing disc tissue mechanics during tissue repair using a mechanical model^10,11^, with a strong purse string healing in the plane of the tissue (*XY*), and lateral cables reducing the tissue height (*Z* axis) to achieve baseline healing dynamics. Here, we utilise our 3D vertex model to modify the initial resting geometry of cells in a multicellular tissue to explore the impact of cell geometry on wound healing strategy. With our model, we predict that more squamous tissues would require a stronger purse string to achieve efficient wound healing compared to columnar tissues. We hypothesise the existence of a regime dependent on the cell aspect ratio and size of the wound beyond which tissues favour alternative repair mechanisms instead of a contractile purse string.

## Methods

### Aspect ratio of cells

We have measured the aspect ratio (*AR*) of cells in different published works (**Table S1**) as follows:

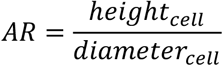

where we measure the diameter of the cell in the apical or basal domain (*XY* plane) assuming *diameter*_*apical*_ = *diameter*_*basal*_ . The height of the cell reflects the length in the apico-basal (*Z*) axis.

### 3D vertex model

We have used our recently developed 3D vertex model^10^. Briefly, the model computes at each time-step *t*_*k*_ the vertex positions 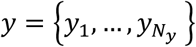 by minimising the total energy of the system *W*. This energy is the result of different contributions:

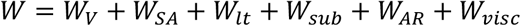

where *W*_*V*_, *W*_*SA*_, *W*_*lt*_, *W*_*sub*_, *W*_*AR*_, *W*_*visc*_ stand respectively for volumetric, surface area, line tension, substrate, triangles’ aspect ratio and viscous energy terms. They are defined by:

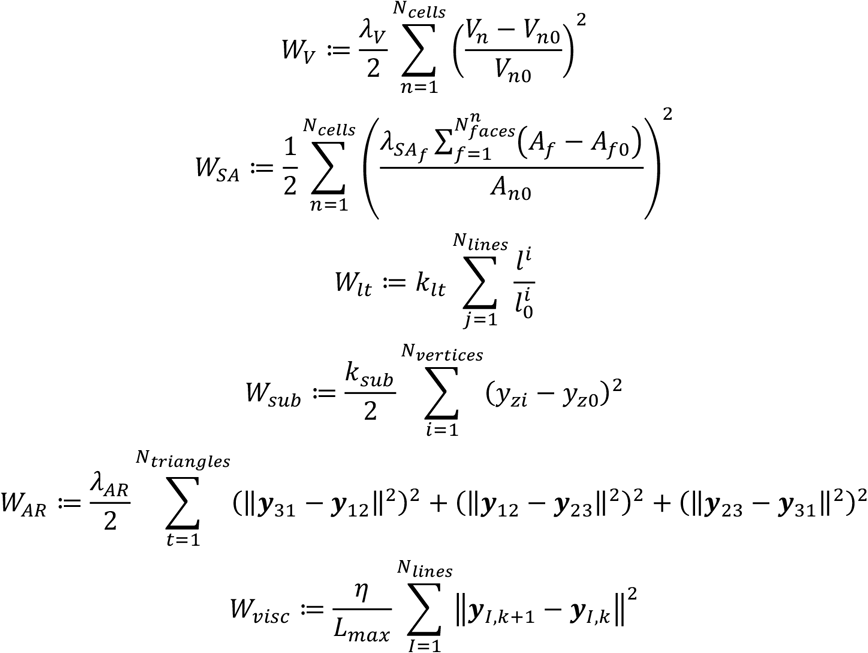

with *l*_*j*_ is the length of line segment *j*, ***y***_*J,0*_ contains the coordinates of vertex *I* at time *t*_*k*_, and 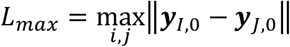. The weighting parameters 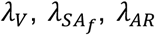, and *η* have been initially calibrated according to recoil evolution in the *Drosophila* wing disc^10^. Further fine-tuning of the parameters is described in the Results section, for matching three-dimensional aspects and for differentiating different domains of the tissue, and during wound healing.

The coordinates ***y***_*I*_ of each vertex *I* are found by computing the positions that minimise the total energy *W*, that is, by solving the following set of equations,

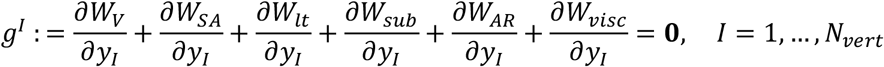

The coordinates of the vertices are updated following a forward explicit Euler solver,

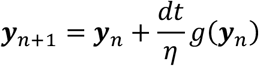

The constant *η >* 0 is the friction term, and *dt* is the current time step. More information can be found in the methods section and supplemental information of our previous manuscript^10^. To account for tissue tension, we have used the contribution from the surface area of the cell that can create tense or loose tissue states. Therefore, we have not used any line tension contribution in a homeostatic state. The line tension contribution (*W*_*lt*_) becomes important for line elements like the contractile purse string, appearing only at the wound edge cells (and only at the line shared between a debris cell and an alive cell). Given that 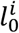 would scale with tissue aspect ratio, and to faithfully compare the purse string strength in different cell aspect ratios, we modelled the line tension of the purse string *W*_*lt*_ with 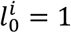 in all cases. Thus, effectively:

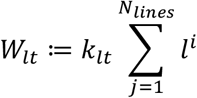

### Geometry

To use realistic cellular topologies, we have used 10 different topologies from the apical domain of the wing disc in larval or pupal stage ^31^. First, we created the different 3D geometries for each topology with an aspect ratio of 15 (like the *Drosophila* wing disc) using the segmented images (**Fig. S1B**). Then, we used the relationship of the diameter of the cell and the cell height to calculate each desired aspect ratio (**Fig. S1A,C**), *AR* = {0.0015, 0.015, 0.15, 1.5, 7.5, 15, 30}. Then, we scaled each in silico tissue such that the volume difference between the reference (*AR* = 15.0) and all the tissues with the same topology is < 1*e* − 19.

Note that all tissues have periodic boundaries, meaning that all vertices (including boundary cells) are connected to at least three other vertices.

### Cell shape analysis with a 3D vertex model and Optuna

We have used Optuna^32^ to obtain what are the parameters that keep the geometry without notable changes. We used a simplified version of our equations, where we only have surface area minimisation and volume preservation:

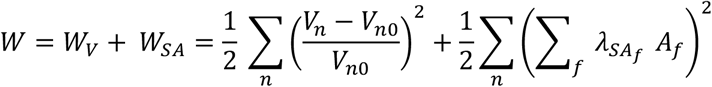

With *λ*_*V*_ = 1, and no surface area elasticity to minimise bias of the different aspect ratios initial surfaces. With Optuna we aimed to find the surface area parameters (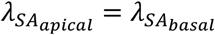, and 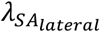) that minimise the energy function (*W*). Those would give us an understanding on which mechanical parameters are allowed to maintain cell shape. This can also be reproduced using the napari plugin napari-pyVertexModel by clicking ‘Estimate *λ* S parameters’ for a given cell height and tissue topology (**Fig. S2**).

### Wound healing mechanics

Prior to all wound healing essays, we performed a series of steps to obtain a pre-equilibrated in silico tissue.

#### Cell(s) ablation

We can ‘kill’ a different number of cells in the middle of the tissue. Ablated cells become ‘debris’ cells that differ from regular cells in their contribution to the energy. Debris cells do only contribute to *W*_*v*_ with a *λ*_*v*_ *= 1e − 8* (regular cells are *λ*_*v*_ *= 1*). To assess the wound recoil at a given *t* time, we compute:

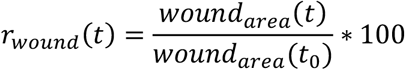

where *wound*_*area*_ is the total surface (typically at the apical side) of all the debris cells. This equation was used to compute ‘apical max recoiling’ of a wound.

#### Edge ablation

To perform an edge ablation, we remove the surface tension on that edge 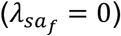. We always perform edge ablations on the apical side since wound healing is driven by the apical side during the first 60 minutes in the *Drosophila* wing disc^11^. To quantify its recoil, we compute how the perimeter of the edge evolves over time, measuring the time evolution of the following non-dimensional quantity, defined by:

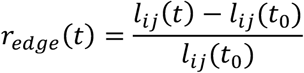

with *l*_*ij*_ is the edge connecting the vertices *i* and *j* whose length is |***y***_*i*_ *−* ***y***_*j*_|, at time *t* and *t*_*0*_ the initial time.

#### Wound closure rate

We defined wound closure rate as the max wound area at 6 minutes (peak recoiling time) minus the final wound area divided by the last timepoint:

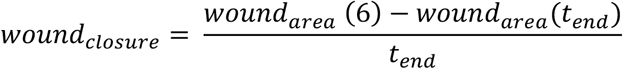

where *t*_*end*_ is 30 minutes or the timepoint the simulation ended.

Finally, to model the purse string for 30 minutes post-ablation, the purse string strength followed the same trend as the *Drosophila* wing disc ^11^, where it begins to be active at approximately 6 minutes.

### Purse string strength required analysis

To compute the purse string strength required to close a gap of a given number of cells, we created a wound by in silico ablating those cells. We let it recoil for a very short time from *t* = 0 to *t* = 0.1. Then, we used the wound area as a reference. We tested different 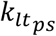 strength 1000 values linearly spaced from 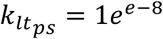 to 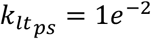. We computed the minimum 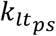 so that the wound area obtained is greater than the initial wound area. Note that *dt = 1e*^−10^.

### Statistical fittings

To understand how features like purse string strength or mechanical parameters like 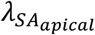, evolves over the aspect ratio, we fitted a function using scipy’s curvefit^33^ that uses non-linear least squares.

### Code availability

The code is available at https://github.com/Pablo1990/pyVertexModel. Instructions on how to run the code are also in the repository. If you want to use the model, you can also use the related version for napari: https://github.com/Pablo1990/napari-pyvertexmodel with Graphical User Interface (GUI) as **Fig. S2**. Installation details can be found there.

### Data availability

All data to reproduce this study is located at the following *Zenodo* link: https://doi.org/10.5281/zenodo.20798921.

## Results

### Differential cell mechanics are defined by the cell’s aspect ratio

We collated a table with studies performing small wounds to epithelial tissues related to the aspect ratio of the cells (**Table S1**). We defined the aspect ratio as cell height divided by the apical or basal cell diameter (see **Methods**). Based on the aspect ratio, we can see a trend where tissues composed of flatter cells do not use purse string as the main healing mechanism (**Table S1**). For instance, the epithelium covering the upper surface of the *Clytiamedusa* called exumbrella is a monolayer of transparent, squamous epithelial cells that are approximately 50μm wide by 1-2μm height (aspect ratio = 0.03)^25^. Their default mechanism is to crawl, but in absence of extracellular matrix (ECM), they sometimes display a purse string driving healing. Likewise, extremely thin cells like corneocytes or alveolar type I pneumocyte (lungs), with aspect ratios close to 0.001, are incapable of migrating or creating other actomyosin-based mechanisms, being unable to effectively repair. Therefore, death, proliferation or differentiation are used by this tissue to heal itself ^34^. In contrast, cuboidal and columnar cells do use a ‘strong’ purse string as the main driver to close wounds in tissues (**Table S1**). For example, in the *Drosophila* 3^rd^ instar larvae wing disc with an aspect ratio of 15, the apical domain leads the healing process with a strong purse string^11^. Similarly, in two different stages of the *Drosophila* embryo with two different cell aspect ratios (stages 7-8: 5.4 aspect ratio, and stages 14-15: 0.5 aspect ratio), it was shown that they use different ways of healing: the *Drosophila* embryo stage 14-15 epidermis exhibits a purse string with filopodia that extends into the wound; while *Drosophila* embryos stage 7-8 that are more columnar appeared to assemble actin protrusions in wounded cells^35^. Other tissues like the zebrafish tail fins epidermis^36^, the wing of the *Drosophila* pupae^37^ and *Xenopus* embryo also seem to use purse string as a healing mechanism, but in the latter, it is thought to be a passive element^8^. However, it is very challenging to compare the real strength of the purse string with regard to the tissue aspect ratio, considering that they have different mechanical environments, are from different organisms, and measured with different techniques^38^.

Thus, we have tackled the importance of the purse string from a theoretical point of view. We developed a computational 3D vertex model where we can ablate cells in silico^10^ and create small wounds in tissues with different shapes (**Fig. S1**). To faithfully compare different tissue shapes, we maintained the initial tissue and cell volumes but changed the cell height (or tissue thickness). Therefore, cells changed their aspect ratio while maintaining their volume, allowing us to compare the different mechanical parameters. To assess the role of cell aspect ratio in determining surface tension, we simplified our model to two energy terms: the bulk of the cell (*W*_*V*_) and its surface tension (*W*_*SA*_),

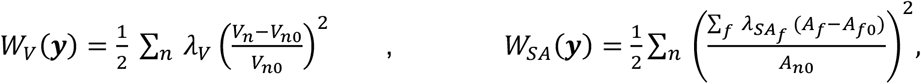

where *A*_*n0*_ *= 1* and *A*_*f0*_ *= 0*, effectively as a purely elastic tensile element. In the same way, we modelled the bulk of the cell as an elastic element with *λ*_*V*_ *= 1*, and *V*_*n0*_ *= V*_*n*_*(t*_*0*_), i.e. the volume at the initial time. Thus,

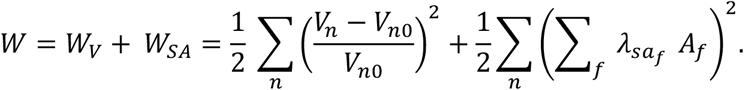

Using this simplified model and the Optuna framework^32^, we looked for the surface area parameters in the different domains 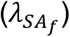 that maintained the aspect ratio of the cells. Epithelial cells typically display an apico-basal polarity, meaning that there are three different surface area domains: apical, basal, and lateral, with different weights in the surface energy, *λ*_*s1*_, *λ*_*s2*_, and *λ*_*s3*_, respectively. We assumed that the apical and basal domains have the same parameter value (*λ*_*s3*_ *= λ*_*s1*_), given that we are only changing cell height and cell aspect ratio. To elucidate how the lateral (*λ*_*s2*_) and apical and basal (*λ*_*s3*_ *= λ*_*s1*_) domains are related, we normalised them for each aspect ratio (**Fig. 1A-A’**). We observed that when cells grow in height (*AR >* 0.15), stronger forces are required in the apical and basal faces to maintain cell shape compared to their lateral side, whereas more isotropic forces are required when cells are more squamous (*AR* < 1.5). The allowed maximum contribution of *λ*_*s2*_ seems to start at a minimum at a 50% when cells are very squamous. Consequently, to quantify the relation between forces in the apical and basal domains and the lateral side, we can fit a sigmoid function, typically used for resource sharing, to understand how the ponderation of the different domains varies (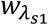 and 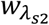) with cell aspect ratio (*x*):

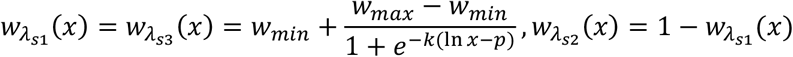

being *x* the aspect ratio of cells, *p* = *ln(x*_*0*_) locates the midpoint, *k* controls steepness, and *w*_*min*_ and *w*_*max*_ sets the lower and upper limit of 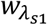, respectively. After fitting with our data, we obtained

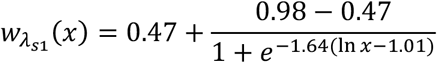

confirming the minimum of *λ*_*s3*_ *= λ*_*s1*_ at around of 50% and upper limit of approximately 1, proposing that infinitely tall cells would need an almost negligible contribution from the lateral surface. In addition, we observed a high variability with a very small aspect ratio (*x* = 0.0015, **Fig. 1A-A’**), implying that the surface area alone cannot control 3D cell shape in this aspect ratio.

**Figure 1.**
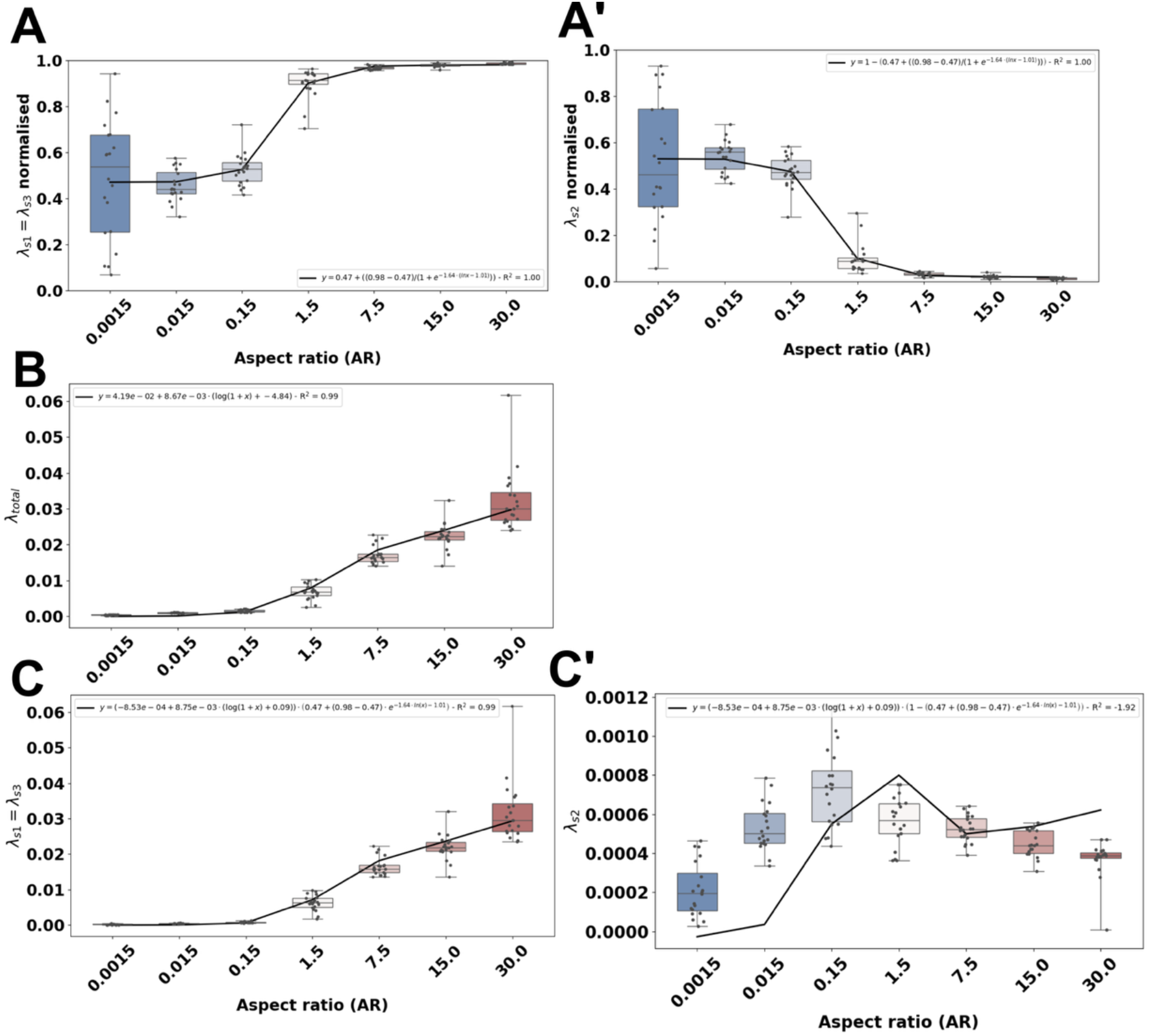
Cell aspect ratio influences the permitted surface tension in different domains. A) Fraction of each surface area parameters **λ**_**s1**_ (apical) and **λ**_**s3**_ (basal), **λ**_**s2**_ is lateral regarding the different aspect ratios. B) **λ**_**total**_ = **λ**_**s1**−**3**_ + **λ**_**s2**_. C-C’) Final fitting to **λ**_**s1**_ = **λ**_**s3**_ and **λ**_**s2**_. Note that colours in panels are related to the aspect ratio (AR). Dark blue: 0.0015. Blue: 0.015. Light blue: 0.15. White: 1.5. Light red: 7.5. Red: 15.0. Dark red: 30.0.

Given a finite quantity of contractile machinery (e.g., myosin II, actin, …) that can be recruited by the apico-basal and lateral domains, we assumed there is a single pool (*g*) that might be changing to regulate cell aspect ratio (*x*) in a short period of time. Thus, we considered:

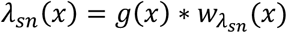

with *n* ∈ {1, 2, 3}, and *g*(*x*) being the contractility machinery pool that depends on the aspect ratio of the cell (*x*). To determine *g*(*x*) behaviour, we computed *λ*_*total*_ *= λ*_*s1*_ *+ λ*_*s2*_ (**Fig. 1B**). Interestingly, we observed that in squamous cells the contribution of the surface area is required to be small (*x* ≤ 0.15). In contrast, from cuboidal aspect ratios (*x* ≥ 1.5), *λ*_*total*_ increases with cell aspect ratio. This possibly means that squamous cells may rely more on volume than surface area. Then, we fitted a log-scaled parabola to the data (**Fig. 1B**), so that:

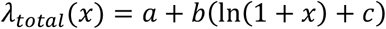

with *a* representing the minimum pool of contractile machinery, *b* how steep is the parabola, and *c* is the aspect ratio representing the minimum *λ*_*total*_. After fitting *λ*_*total*_, we got *a* = 4.19*e*^− 2^, *b* = 8.67 *e*^*–3*^, and *c* = −4.84 (**Fig. 1B**). By combining both functions of *λ*_*total*_ (*x*) representing the total amount of contractile machinery, and 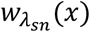 how the machinery will be distributed between the apico-basal and lateral domains, we obtained the final value of *λ*_*sn*_ is:

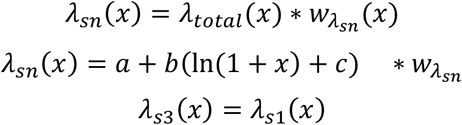

As predicted (**Fig. 1A-A’**), from cuboidal to columnar aspect ratios, the apico-basal surface area strength should increase to maintain cell aspect ratio, implying an increment in the quantity of contractile machinery (**Fig. 1B-C**). Remarkably, whereas *λ*_*s1*_ and *λ*_*s3*_ grow linearly with cell aspect ratio, *λ*_*s2*_ peaks at an aspect ratio of 0.15 (**Fig. 1C’**), which may be related to the average number of lateral faces ⟨*n*_*faces*_⟩ *≈ 6* with *1/*⟨*n*_*faces*_⟩ *≈ 0*.*15* (see **Discussion**).

Overall, in cells with small aspect ratios, the surface tension needs to be isotropic to maintain the cell’s shape. In contrast, when cells have bigger aspect ratios (*x* ≥ 1.5), the anisotropy between the apico-basal domain and the lateral side grows, consistent with other studies^39^. Given that we see a decrease in surface tension in the apical domain when cells are flatter, would the tissue be able to use the same wound healing mechanisms to heal optimally, independently of the aspect ratio of the cells?

### 3D cell shape affects healing mechanics

In our previous work, we acquired the mechanical parameters to obtain the *Drosophila* wing disc during homeostasis and wound healing^10^, where a contractile purse string drives the healing of the wound. Previously, we modelled the purse string as contractile line elements or line tension:

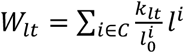

where *k*_*lt*_ controls the tensile strength, *l*^*i*^ is the length of the edge *i* and 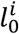 its reference length. Note that this contractility will only be used for the purse string, the resting tension derives from the surface tension. Also, to faithfully compare different structures, we modelled the purse string as a purely elastic element with 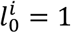. Further details in **Methods**. To obtain a biologically relevant surface tension, we used *λ*_*s*1_= 1.4 and a *λ*_*s*2_ = 0.014 to mimic *ex vivo* results, including laser ablation-induced wound recoiling profiles^10^, to close a gap of 10 cells as performed *ex vivo* in *Drosophila*. To meaningfully compare all the different aspect ratios, we use the apical edge recoil to compare the tension in tissues with different aspect ratios. Here, the edge recoil is a proxy for the effective resting surface tension. Thus, we obtained simulations with a similar apical edge recoil (**Fig. 2B**). This is set as the physiologically comparable resting tension across the different cell aspect ratios, despite their different absolute mechanical parameters (**Fig. 2B’**). Interestingly, to obtain a given recoil, more squamous tissues require weaker surface tension, consistent with **Fig. 1**, where taller cells are more energetically stable with stronger apical and basal surface tension (**Fig. 2B-B’**).

**Figure 2.**
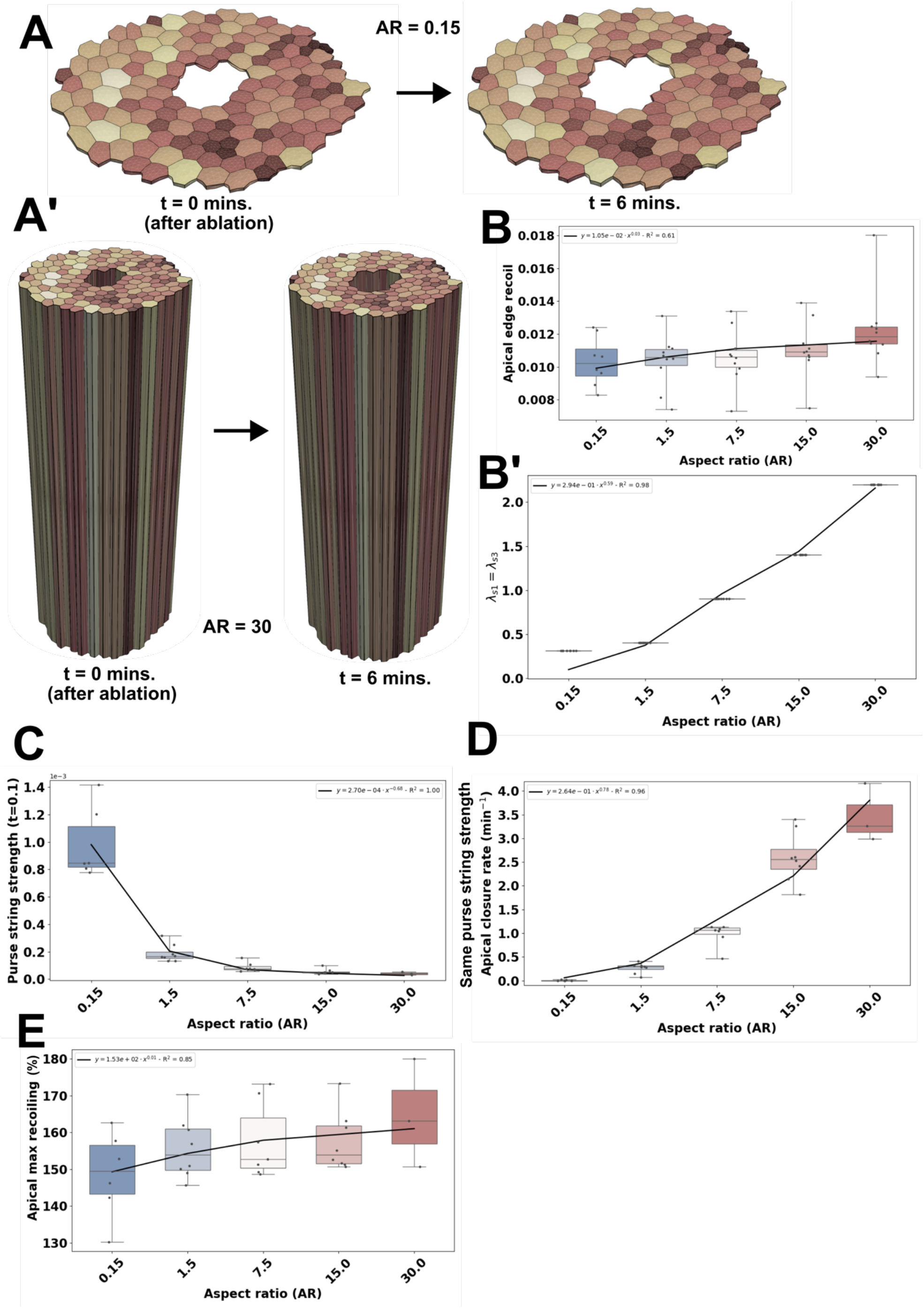
Despite having the same wound recoil, flatter tissues require a stronger purse string. A) Illustrative figure of simulations with aspect ratios (AR) of 0.15 and 30 (A’) showing ablation and recoil at 6 minutes. Note that volumes or shapes are not preserved in the panel. Colours represent the range of volume using the same limits in both simulations. B) Apical edge recoil (**recoil**_**edge**_, dimensionless) of tissues with different aspect ratios. B’) Shows lambdaS1 to achieve those recoils. C) Purse string strength predicted by the model at t=0.1 minutes after ablation. D) Using the same purse string strength, apical closure rate (1/minutes). E) Apical max recoiling (%) at 6 minutes after wounding. Note that colours in panels are related to the aspect ratio (AR). Light blue: 0.15. White: 1.5. Light red: 7.5. Red: 15.0. Dark red: 30.0. Fittings show equation derived and **R**^**2**^.

Based on these results, we asked whether the mechanical model could predict the purse string strength required 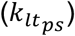 to initiate healing at early stages. Therefore, we computed the minimum purse string strength needed to close the gap right after ablation (*t* = 0.1 minutes). Indeed, the model predicts that a purse string 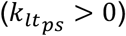 is needed for the wound to start to close, regardless of the aspect ratio (**Fig. 2C**). Furthermore, the minimum purse string strength needed to close the wound decreases as the aspect ratio of cells increases (**Fig. 2C**). In other words, the biophysical model suggests that, eventually, the purse string alone would not be able to heal very flat cells, because a close-to-infinite force would be required. This is consistent with the experimental observation that very flat types of cells are unable to heal^34^. To quantitatively assess the model predictions, we run the model for 30 minutes after ablation with a fixed purse string strength 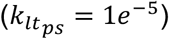 for the different cell aspect ratios (**Movie S1-5**). Then, at 30 minutes, we analysed how much area it has closed with regard to its maximum recoil area and relative to time, namely the apical closure rate. Consistent with our previous predictions (**Fig. 2C**), we can see taller cells would heal very efficiently, whereas squamous cells will not heal at all meaning that the tissue will recoil and never close the wound (**Fig. 2D**). This confirmed our previous predictions and supports this simple but not straightforward idea that, given the same cell volume, shorter tissues would require a stronger purse string than taller cells to close a wound. However, given that we have ablated the same number of cells in all cases, does the number of ablated cells impact tissue repair?

### Ablating the same wound area in different tissue aspect ratios

A common question that remains unanswered in the field is whether we can infer the healing mechanism of a tissue by ablating the same absolute surface area. While it has been seen that on *in vitro* adhered tissues, wound area plays a role in the closure rate^40^, it is unknown how ablating different surface areas in tissues will affect the healing mechanism *in vivo*. With our computational model, we have undertaken this question from a biophysical viewpoint. We used the same simulations as before (**Fig. 2**), where we have the same edge recoil, and, therefore, same tension regardless of geometry. Instead of ablating 10 cells, we computed the number of cells needed to create a gap with the same absolute *X*-*Y* area. We used our smallest aspect ratio that is able to heal (*AR* = 0.15) as our baseline, where we ablated a single cell. Then, for *AR* = 1.5, we needed 5 cells, for *AR* = 7.5, 14 cells, for *AR* = 15, 23 cells, and *AR* = 30, 35 cells, respectively, to replicate the same wound area (**Fig. 3A**, and **Fig. S4**). Interestingly, the wound recoiling profiles are very distinct (**Fig. 3B**, and **Fig. 2E**), even though the edge recoil is the same among different samples (**Fig. 2B**). Smaller wounds in shorter tissues had a strong wound recoil, whereas bigger wounds in taller tissues had a weaker recoil. This counterintuitive prediction suggests that the number of ablated cells affects wound healing dynamics. We performed the same experiment as **Fig. 2D** where we used the same purse string strength 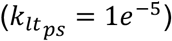, and found that, even though the recoiling area percentage was not massively affected (except in squamous tissues), their healing ability was impacted. All aspect ratios displayed a similar (*AR* ≤ 1.5) or delayed (*AR >* 1.5) apical closure rate compared to a 10 cells ablation (**Fig. 3C**, and **Movie S6-10**). The model also shows that the number of ablated cells influences the required purse string strength to initiate closure (**Fig. 3D**). This may be due to how we have modelled the purse string as a collection of linear rods rather than a continuous exponential string (see **Discussion**). Overall, these results imply that the force required to initiate gap closure is different from healing a wound completely. The former relates to the initial ablated state (e.g., numbers of ablated cells, resting line tension), whereas the latter considers the total area to close over time, where the aspect ratio of tissue is predominant (**Fig. 2C-D** and **Fig. 3D-E**). Finally, given that we saw a change in apical closure rate when ablating different number of cells, we performed simulations that ablated a single cell (**Movie S6**, and **Movie S11-14**). This resulted in the same paradigm shown before: the apical closure rate follows an exponential decay when approaching small aspect ratios (**Fig. S4**). In addition, the computational model predicted that ablating a smaller subset of cells would result in faster healing (**Fig. 3E**). Still, regardless of the recoil and the number of ablated cells, flatter tissues (*AR* ≤ 1.5), exhibited poor healing rates. For more columnar cells (*AR >* 1.5), we saw that ablating a single cell made the tissue heal at a faster rate, achieving full closure in less than 30 minutes (**Fig. 3F**), even though the wound recoil is larger (**Fig. 3G**). In contrast, ablating 10 or more cells resulted in a slower healing rate and even healed poorly (**Fig. 3E**). Thus, implying that smaller wounds would require a weaker purse string than bigger ones. This is consistent with previous studies showing that the purse string is only effective in small wound sizes (see **Discussion**).

**Figure 3.**
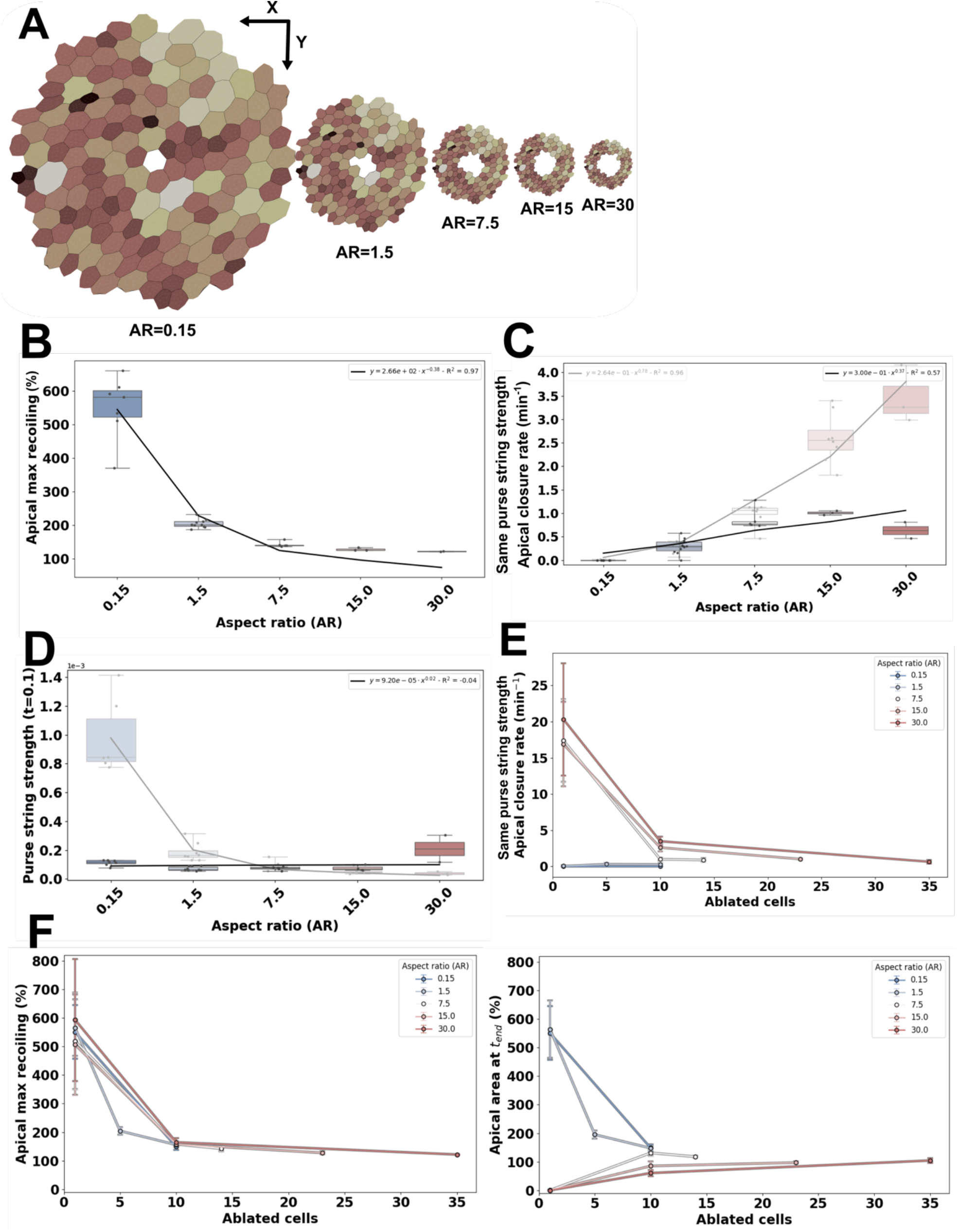
Stronger purse strings are needed when more cells are ablated. A) In silico tissues displaying different aspect ratios and different number of ablated cells, but the same wound area. B) The apical maximum recoiling percentages related to initial wound area for each aspect ratio. C) Apical wound closure rate using the same apical purse string strength parameter. Shaded boxplot relates to Fig. 2D. D) Purse string strength parameter required to initiate wound closure at t=0.1 after ablation. Shaded boxplot refers to Fig. 2D. E) Apical wound closure rate regarding number of ablated cells. F) Left, Apical max recoiling (%) and, right, apical area (%) at final timepoint regarding the number of ablated cells. For E and F, points are mean and error bars are standard deviation. Note that colours in panels are related to the aspect ratio (AR). Light blue: 0.15. White: 1.5. Light red: 7.5. Red: 15.0. Dark red: 30.0. Fittings show equation derived and **R**^**2**^.

## Discussion

The so-called purse string, a contractile ring that appears in small wounds, is a mechanism by which epithelial tissues aim to restore homeostasis by closing any possible gap. However, previous works have shown the purse string appearing in a range of cell aspect ratios with varying results. Here, we have performed 3D in silico experiments to better understand why the purse string appears effective in some circumstances. We hypothesise that the purse string would be more efficient in columnar tissues compared to squamous cells as our biophysical model predicts. This would explain why squamous epithelia would only use migration or a partially effective purse string to repair themselves^4,8^. In contrast, columnar cells can rely on the purse string as the main driver^5,11,41^.

Our 3D model predicts that as cells become more columnar (with a higher aspect ratio), the contractile machinery at the apical and basal domains, such as myosin II or actin filaments, may increase to help maintain cell shape. However, this additional recruitment would not be required in squamous (flatter) cells (**Fig. 1**). In contrast, the lateral domain peaks at around 0.15, which could be approximated to the average number of lateral faces (or 3D neighbours) ⟨*n*_*faces*_⟩ ≈ 6 with 1/⟨*n*_*faces*_⟩ ≈ 0.15. Previous work suggested that the number of 3D neighbours may affect the energy landscape of tissues^42^, implying that the number of spatial intercalations may change the interplay between the lateral, and apical and basal domains. To model the purse string as a contractile cable, we have used a purely elastic equation whose energy only depends on the edge’s current length. In particular, we modelled the purse string as the sum of the contributions of the cells’ edges at the wound border. In this context, given the expression of the contractility equation and the linearity of the line tension energy, the number of cells does not have any explicit effect on the final strength: 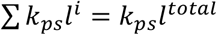. However, non-linear constitutive laws may introduce some additional dependences on the number of segments than those considered here.

As seen in **Fig. 2** and **Fig. 3**, in squamous tissues, even a tiny wound of a single cell would require very strong purse string forces to close it. This theoretically makes the purse string an unlikely mechanism to be used by flatter cells. This is consistent with previous works showing that cells with very flat extreme aspect ratios, which can’t divide (e.g. post-mitotic), will not be able to heal via purse string^4,34^. In contrast to other biological models (like *Drosophila*), other factors, like the role of the substrate, may explain why the purse string is suggested not to be functional in Xenopus ectoderm pigmented cells (superficial layer), despite exhibiting a cuboidal aspect ratio and an apparent purse string^8^. Our model does not also predict why very tall cells, such as fibre-like cells like neural cells, do not employ a contractile cable to seal gaps and rely on other mechanisms to heal, like calcium waves^43^ or other cells like Schwann cells^44^.

A possible physical explanation for the faster closure rates in epithelia with larger aspect ratios *AR* (**Fig. 2**, and **Fig. 3**) may be found in the role of the lateral areas as an energy buffer. Indeed, the surface area ratio between the lateral (*S*_*L*_) and apico-basal areas (*S*_*AB*_) can be expressed as a function of the aspect ratio as *AR* = 0.5*S*_*L*_/*S*_*AB*_, after approximating the cell by a cylinder whose volume is 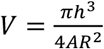. Therefore, larger values of *AR* involve larger ratios of *S* /*S*, and a dramatic increase in the height *h*, since the cube of the cell’s height (*h*^3^) scales with the aspect ratio of the cell squared (*AR*^5^) times *V*, with *V* the cell volume, assumed constant. We consequently hypothesise that the lateral surfaces may function as energy sources for the purse string mechanism. Biologically, this could be represented by a membrane reservoir aiding the cell to accommodate its deformation without increasing membrane tension, like the work by Kosmalska & cols^45^. Here, they showed that the membrane of the cells can store fractions of its surface area as a membrane reservoir. In this context, columnar cells can use their lateral domain as this reservoir. This is also coherent with its a low-tension regime shown in **Fig. 1**. Squamous cells, lacking this buffer, would be penalised for changing shape in this way, thereby slowing down the healing process or necessitating alternative methods to the purse string.

In the same way, results are in concordance with the purse string not being used in larger wounds. As seen in **Fig. 3**, when we ablate more cells, the model predicts that a stronger purse string is required, meaning that large wounds would require too much contractile machinery to be recruited to form an extremely strong purse string. This agrees with previous works with non-adhesive gaps^46,47^, suggesting that there is a limit in gap size in which a contractile ring is no longer able to close the gap. We also suggest that the 3D shape of the gap is important to choose the closure mechanism, particularly in flatter cells.

To sum up, we have used our 3D computational model to theoretically explore the purse string efficiency during wound healing with regard to tissue aspect ratio. We foresee that our results will guide new venues to be tested experimentally in vivo in different organisms. This will be needed to explain how the shape of an organ affects its ability to heal and why the geometry and the mechanical environment affect the chosen mechanism for gap closure. In the long term, our biophysical model could aid personalised medicine in creating realistic digital twins of cancer models and beyond.

## Acknowledgments

We thank the Mao lab for comments on our work, particularly, Veronika Lachina and Giulia Paci for providing feedback on the manuscript. P.V.-M. was supported by EPSRC grant EP/X03139X/1. Y.M. was supported by the MRC award MR/W027437/1, a Lister Institute Research Prize and EMBO Young Investigator Programme.

## Supplementary Tables

***Table S1.**Table with tissues found in literature with the following information: organism, Tissue/Epithelium, purse string (if it was found yes/no), reference for wound healing evidence, aspect ratio calculated from apical or basal cell diameter in micrometres, and cell height in micrometres, reference for the aspect ratio measurements.*

## Supplementary Figures

**Figure S1.**
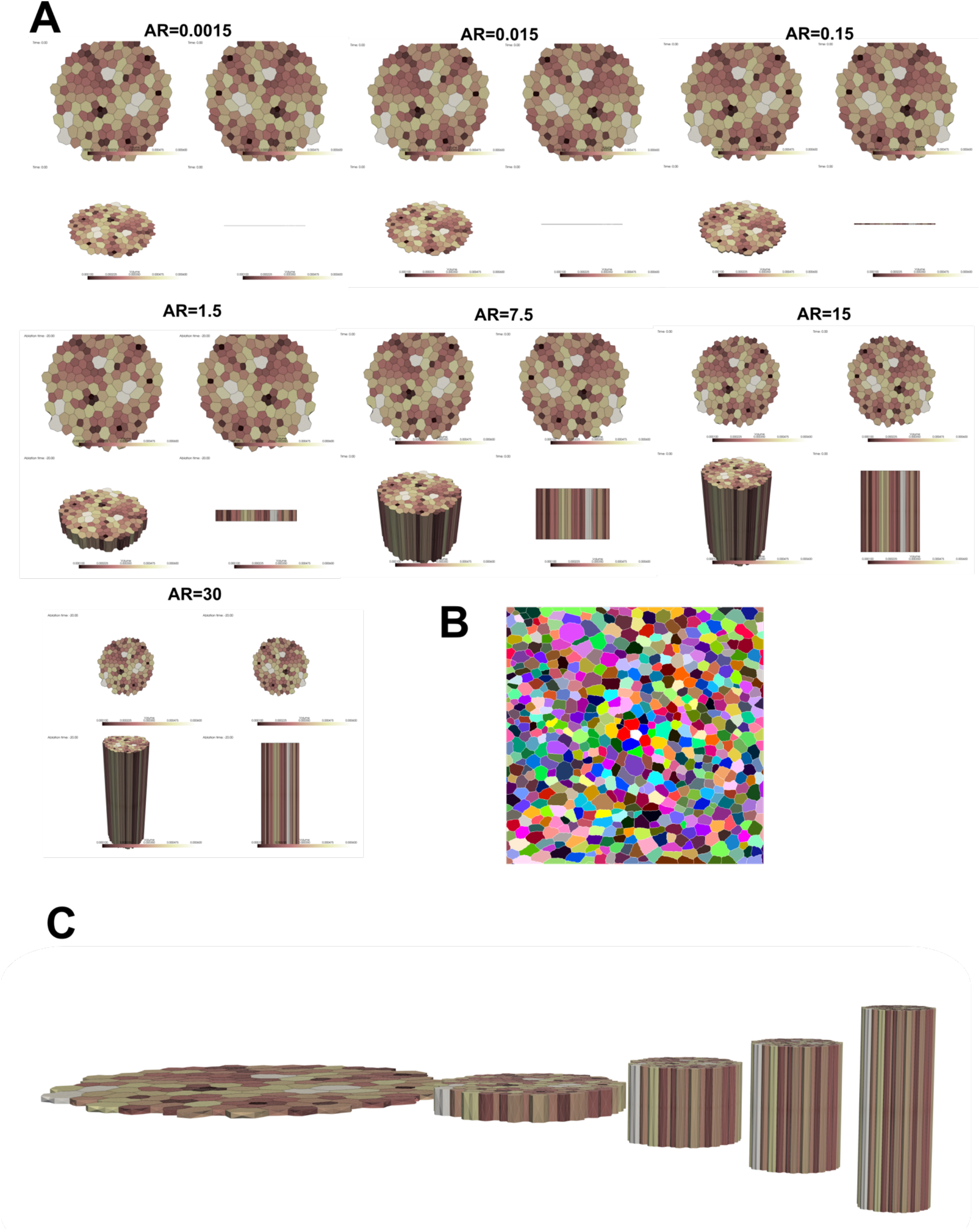
3D topologies used for this paper with images. A) Different views for the different aspect ratios used with the same topology. B) Topology initial image. C) 3D visualisation of the different aspect ratios (from 0.15 to 30).

**Figure S2.**
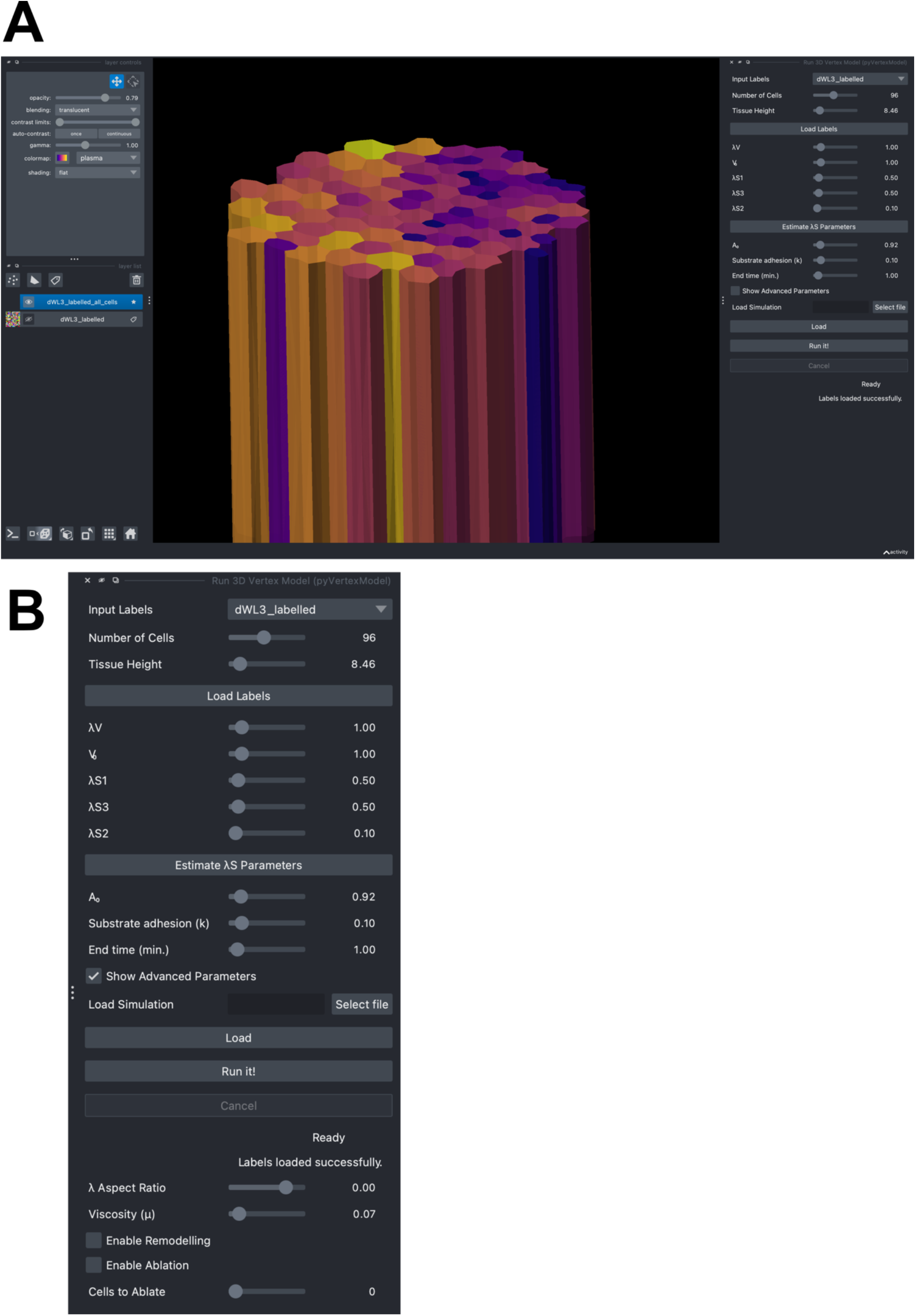
Graphical user interface of napari-pyVertexModel. A) Example of tissue geometry displayed in the napari viewer. B) Different input to the vertex model, buttons, and advanced parameters are not shown by default.

**Figure S3.**
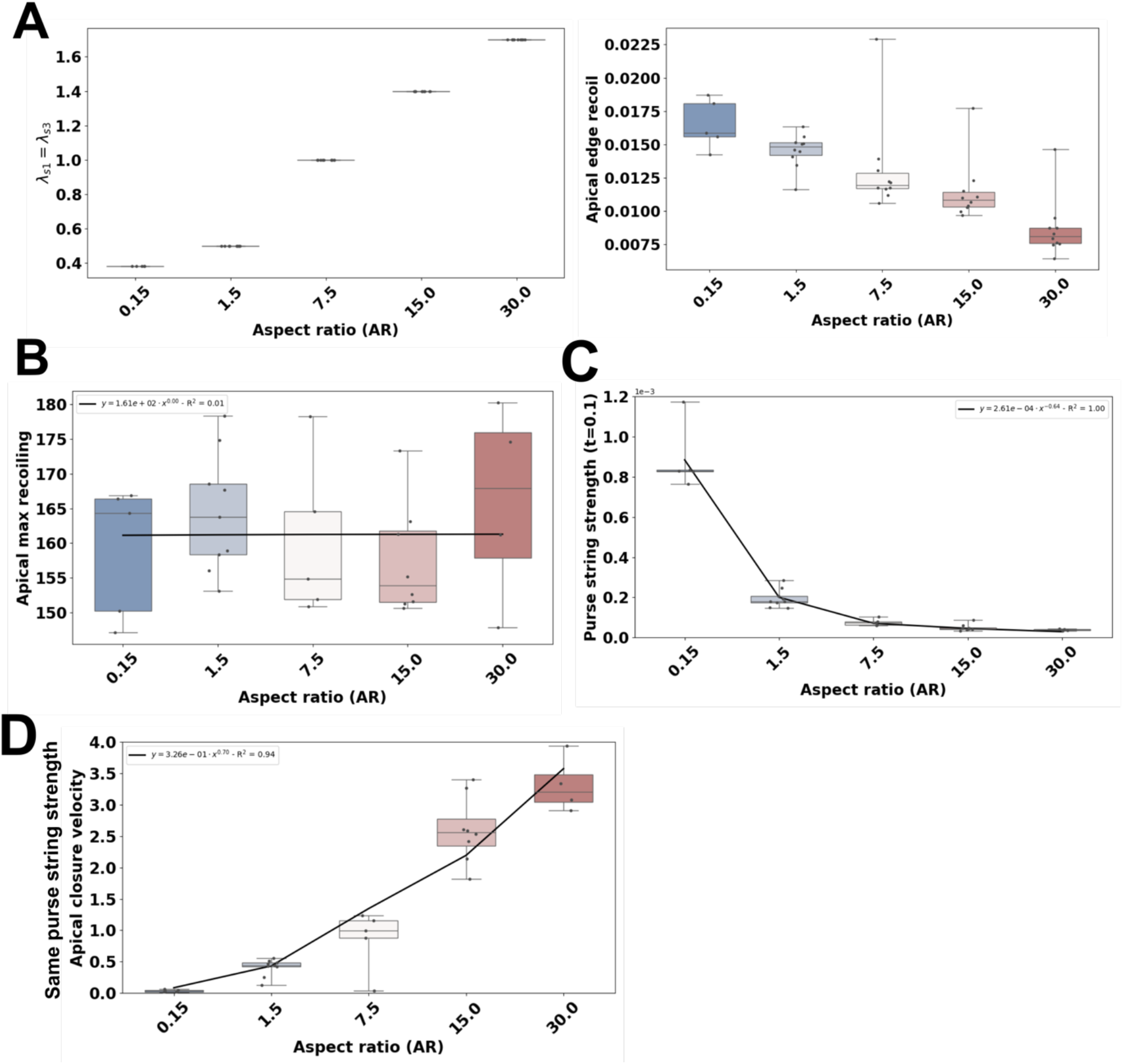
Results with the same wound (10 cells) recoil when 10 cells are ablated, related to Fig. 2. A) Left refers to the \lambda_s1 parameter used in Fig. 3. Right refers to the apical edge recoil of all the simulations. B) Apical max recoiling (%) at t=6 min. C) Purse string strength to initiate wound healing at t=0.1 minutes. D) Apical closure rate (min^-1^) using the same purse string strength. Note that colours in panels are related to the aspect ratio (AR). Light blue: 0.15. White: 1.5. Light red: 7.5. Red: 15.0. Dark red: 30.0. Fittings show equation derived, and **R**^**2**^.

**Figure S4.**
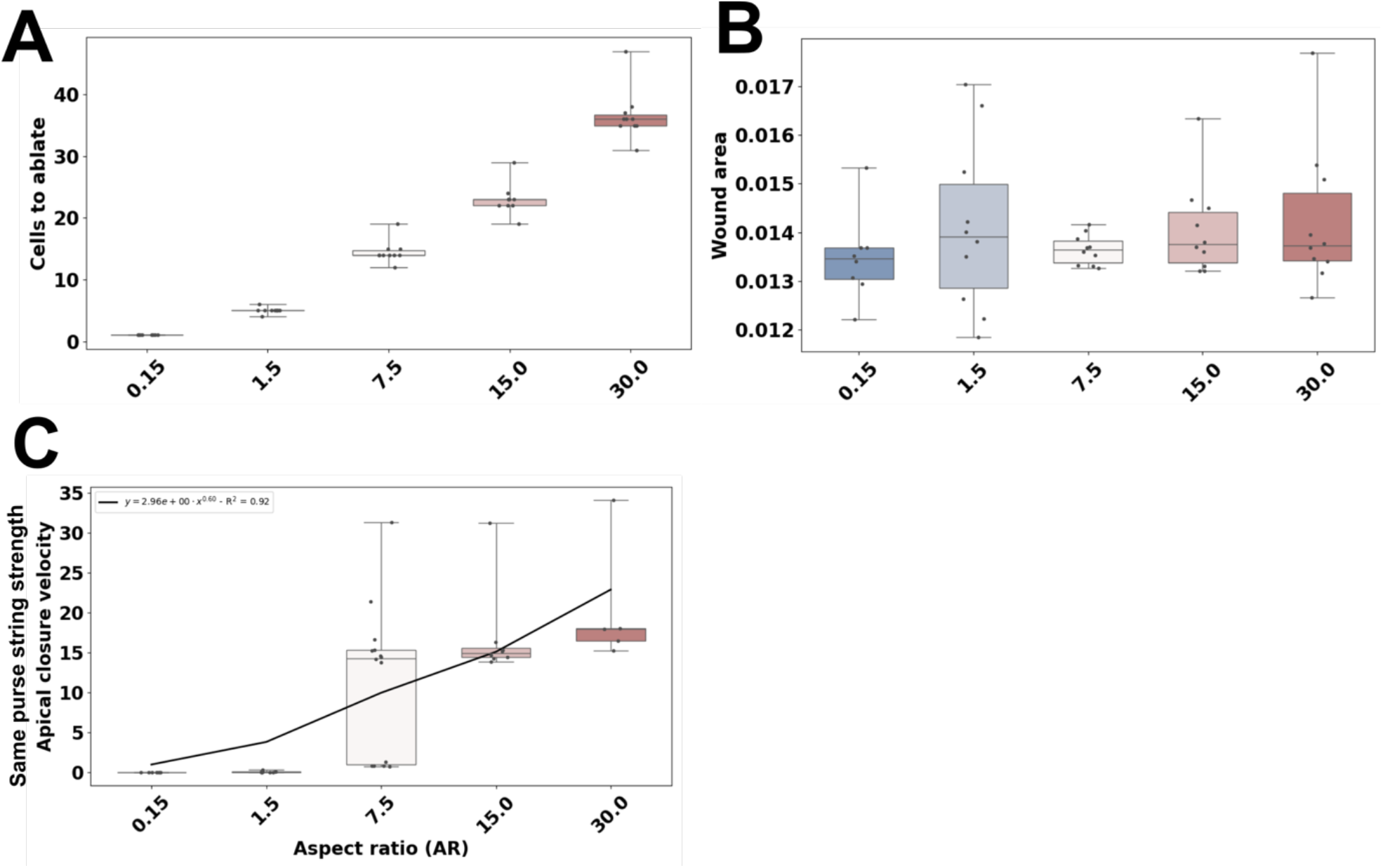
Additional results related to Fig. 3. A) The number of ablated cells on each cell aspect ratio to obtain a similar initial wound area (B). B) The quantified wound initial area when ablating the ablated cells in A. C) Apical closure rate to close a single cell using the same purse string strength. Note that colours in panels are related to the aspect ratio (AR). Light blue: 0.15. White: 1.5. Light red: 7.5. Red: 15.0. Dark red: 30.0. Fittings show equation derived, and **R**^**2**^.

## Supplementary Movies

***Movie S1***. *Biophysical model with aspect ratio 0*.*15 over healing dynamics after a 10-cells ablation. Top left: apical view (from top). Top right: basal view (from bottom). Bottom left: perspective view. Bottom-right: lateral view from the middle of the wound with shaded debris cells. Colour of cells represent their volume over time as present in the legend from dark brown to light yellow*.

***Movie S2***. *Biophysical model with aspect ratio 1*.*5 over healing dynamics after a 10-cells ablation. Top left: apical view (from top). Top right: basal view (from bottom). Bottom left: perspective view. Bottom-right: lateral view from the middle of the wound with shaded debris cells. Colour of cells represent their volume over time as present in the legend from dark brown to light yellow*.

***Movie S3***. Biophysical model with *aspect ratio 7*.*5 over healing dynamics after a 10-cells ablation. Top left: apical view (from top). Top right: basal view (from bottom). Bottom left: perspective view. Bottom-right: lateral view from the middle of the wound with shaded debris cells. Colour of cells represent their volume over time as present in the legend from dark brown to light yellow*.

***Movie S4***. Biophysical model with *aspect ratio 15 over healing dynamics after a 10-cells ablation. Top left: apical view (from top). Top right: basal view (from bottom). Bottom left: perspective view. Bottom-right: lateral view from the middle of the wound with shaded debris cells. Colour of cells represent their volume over time as present in the legend from dark brown to light yellow*.

***Movie S5***. Biophysical model with *aspect ratio 30 over healing dynamics after a 10-cell ablation. Top left: apical view (from top). Top right: basal view (from bottom). Bottom left: perspective view. Bottom-right: lateral view from the middle of the wound with shaded debris cells. Colour of cells represent their volume over time as present in the legend from dark brown to light yellow*.

***Movie S6***. Biophysical model with *aspect ratio 0*.*15 over healing dynamics after a 1-cell ablation. Top left: apical view (from top). Top right: basal view (from bottom). Bottom left: perspective view. Bottom-right: lateral view from the middle of the wound with shaded debris cells. Colour of cells represent their volume over time as present in the legend from dark brown to light yellow*.

***Movie S7***. Biophysical model with *aspect ratio 1*.*5 over healing dynamics after a 5-cells ablation. Top left: apical view (from top). Top right: basal view (from bottom). Bottom left: perspective view. Bottom-right: lateral view from the middle of the wound with shaded debris cells. Colour of cells represent their volume over time as present in the legend from dark brown to light yellow*.

***Movie S8***. Biophysical model with *aspect ratio 7*.*5 over healing dynamics after a 14-cells ablation. Top left: apical view (from top). Top right: basal view (from bottom). Bottom left: perspective view. Bottom-right: lateral view from the middle of the wound with shaded debris cells. Colour of cells represent their volume over time as present in the legend from dark brown to light yellow*.

***Movie S9***. Biophysical model with *aspect ratio 15 over healing dynamics after a 23-cells ablation. Top left: apical view (from top). Top right: basal view (from bottom). Bottom left: perspective view. Bottom-right: lateral view from the middle of the wound with shaded debris cells. Colour of cells represent their volume over time as present in the legend from dark brown to light yellow*.

***Movie S10***. Biophysical model with *aspect ratio 30 over healing dynamics after a 35-cells ablation. Top left: apical view (from top). Top right: basal view (from bottom). Bottom left: perspective view. Bottom-right: lateral view from the middle of the wound with shaded debris cells. Colour of cells represent their volume over time as present in the legend from dark brown to light yellow*.

***Movie S11***. Biophysical model with *aspect ratio 1*.*5 over healing dynamics after a 1-cell ablation. Top left: apical view (from top). Top right: basal view (from bottom). Bottom left: perspective view. Bottom-right: lateral view from the middle of the wound with shaded debris cells. Colour of cells represent their volume over time as present in the legend from dark brown to light yellow*.

***Movie S12***. Biophysical model with *aspect ratio 7*.*5 over healing dynamics after a 1-cell ablation. Top left: apical view (from top). Top right: basal view (from bottom). Bottom left: perspective view. Bottom-right: lateral view from the middle of the wound with shaded debris cells. Colour of cells represent their volume over time as present in the legend from dark brown to light yellow*.

***Movie S13***. Biophysical model with *aspect ratio 15 over healing dynamics after a 1-cell ablation. Top left: apical view (from top). Top right: basal view (from bottom). Bottom left: perspective view. Bottom-right: lateral view from the middle of the wound with shaded debris cells. Colour of cells represent their volume over time as present in the legend from dark brown to light yellow*.

***Movie S14***. Biophysical model with *aspect ratio 30 over healing dynamics after a 1-cell ablation. Top left: apical view (from top). Top right: basal view (from bottom). Bottom left: perspective view. Bottom-right: lateral view from the middle of the wound with shaded debris cells. Colour of cells represent their volume over time as present in the legend from dark brown to light yellow*.

## Notes

### Competing Interest Statement

The authors have declared no competing interest.

